# Dynamic updating of cognitive maps via traces of experience in the subiculum

**DOI:** 10.1101/2025.10.01.679719

**Authors:** Fei Wang, Andrej Bicanski

**Affiliations:** Max-Planck Institute for Human Cognitive and Brain Sciences, Department of Psychology, Leipzig, Germany

**Keywords:** Subiculum, Spatial memory, Cognitive Map, Mismatch, Vector Trace Cells

## Abstract

In the classical view of hippocampal function, the subiculum is assigned the role as the output layer. In spatial paradigms, some subiculum neurons manifest as so-called boundary vector cells (BVCs), firing in response to boundaries at specific allocentric directions and distances. More recently it has been shown that some subiculum BVCs can be classified as vector trace cells (VTCs), which exhibit traces of activity after a boundary/object has been removed. Here we propose a model of processing within subiculum that accounts for VTCs, taking into account proximodistal differences in subiculum (pSub vs dSub) and CA1. dSub neurons receive feedforward input, either in the form of perceptual information (from BVCs in pSub) or mnemonic information (from place cells in CA1). Mismatch between these two inputs updates associative memory encoded in the synapses between CA1 and dSub. With a range of learning rates, the model captures the majority of experimental findings, including the distribution of VTCs along the proximodistal axis, the percentage of VTCs across different cue types, and the hours-long persistence of the vector trace. Incorporating experimentally reported effects of inserted objects/rewards on place cells (place field shift), we also explain why VTCs have longer tuning distances after cue removal. This adds predictive character to subiculum traces and suggests the online use of mnemonic content during navigation. Our model suggests that mismatch detection for updating spatial memory content provides a mechanistic explanation for findings in the CA1-subiculum pathway. This work constitutes the first dedicated circuit-level model of computation within the subiculum, consistent with known effects in CA1, and provides a potential framework to extend the canonical model of hippocampal function with a subiculum component.

## 1 Introduction

The subiculum is commonly viewed as the output layer of the hippocampal formation, receiving direct inputs from the CA1 region and projecting to various cortical and subcortical areas (Amaral and Witter 1989; Matsumoto, Kitanishi, and Mizuseki 2019; Witter 2006). The classical model of the hippocampus suggests that the dentate gyrus performs pattern separation on entorhinal cortex inputs, followed by auto-association in CA3 (via Schaffer collaterals). At the same time, a direct connection from entorhinal cortex forms CA1 inputs, which are then associated with CA3 (hetero-association) (Treves and Rolls 1994). Novel findings and proposals notwithstanding (Rolls and Treves 2024; Stachenfeld, Botvinick, and Gershman 2017), the standard model has strong experimental support. However, the subiculum does not feature in this account.

The subiculum does not simply relay information to downstream brain regions. Recent studies shows that the subiculum holds accurate neural representations of navigation-related variables, such as trajectory and speed (Kitanishi, Umaba, and Mizuseki 2021). Subiculum neurons also exhibit several forms of synaptic plasticity and form recurrent circuits (Berger et al. 1980; Köhler 1985). These network properties imply that subiculum possesses intrinsic computational capabilities. Moreover, contrary to the canonical view as hippocampal output, there is also significant evidence for direct projections from the subiculum to CA1 (Commins, Aggleton, and O’Mara 2002; Jackson et al. 2014; Shao and Dudek 2005; Sun et al. 2014; Sun et al. 2018; Xu et al. 2016), and these projections are of a similar strength as the EC-CA1 projections (Brun et al. 2002). Together with the recurrence in subiculum, this suggests the subiculum may affect the formation and – as we suggest below – the updating of spatial memory and cognitive maps more broadly.

The term cognitive map (O’Keefe and Nadel 1978; Tolman 1948) refers to a neural representation that encodes systematic relationships between elements within an environment to enable flexible behavior. In the spatial domain, it results from associative memory between spatial entities (e.g., landmarks, coordinates, boundaries), helping animals perform navigation tasks (Behrens et al. 2018). Experiments have revealed spatial codes in subiculum (Kim, Ganguli, and Frank 2012; Lever et al. 2009; Sun et al. 2024). Boundary vector cells (BVCs) in the subiculum respond maximally when environmental boundaries are perceived at specific vectors from the animal (Lever et al. 2009; Muessig et al. 2024; Stewart et al. 2014, for review see, Bicanski and Burgess 2020). Experimental recordings of BVCs followed earlier simulation studies, specifically the BVC-to-PC model (Hartley et al. 2000), showing a thresholded sum of postulated BVCs can generate the firing of place cells (PCs). These existing results imply the subiculum’s potential role in supporting associations between scene elements (e.g., boundaries) at specific spatial coordinates.

Subiculum neurons also provide a mnemonic representation of recent experiences (Poulter et al. 2021). A subset of BVCs exhibit persistent firing even after the removal of a previously perceived boundary. This trace phenomenon primarily occurs in the distal subiculum (dSub), whereas most BVCs in the proximal subiculum (pSub) respond only to currently present boundaries. Following the BVC-to-PC model (Hartley et al. 2000), later studies further simulated the imagined BVCs reactivated by PCs in familiar environments, facilitating memory retrieval (Becker and Burgess 2000; Byrne, Becker, and Burgess 2007). Specifically, the so-called BB-model (Bicanski and Burgess 2018) simulated the trace-phenomenon but without distinguishing trace from non-trace cells and without subiculum-internal computations. These models describe the potential functions embedded in the CA1-subiculum circuit, but do not address differences along the proximodistal axis of the subiculum (Poulter et al. 2021). Similarly, differences between proximal and distal CA1 coding have not found their way into models. It has been suggested that proximal CA1 (pCA1) conveys spatial information to the dSub, whereas distal CA1 (dCA1) conveys non-spatial information to the pSub (Igarashi et al. 2014; Kim and Spruston 2012; Matsumoto et al. 2019; Nagelhus et al. 2023; Vandrey, Duncan, and Ainge 2021).

Inspired by the experimental studies that identified vector traces (Poulter et al. 2021), we developed a model of trace coding and the proximodistal differences in the Sub and CA1. We explore the CA1-subiculum network, where dSub neurons receive the inputs from pSub BVCs and CA1 PCs. BVCs in the pSub provide perception information by responding to the present boundaries, including inserted cues (suggested to be derived from upstream sensory processing). The subicular complex—including the subiculum, prosubiculum, presubiculum, postsubiculum, and parasubiculum—has been shown to receive projections from various cortical and subcortical regions, such as the temporal cortex, retrosplenial cortex, and perirhinal cortex (Aggleton 1986; Ding 2013; Kosel, Van Hoesen, and Rosene 1983; Rosenblum et al. 2024; Yukie 2000). PCs in CA1 can drive the transmission of corresponding mnemonic information through CA1-dSub connections, which encode associative spatial memories linking boundaries to spatial coordinates. The firing rate of each dSub neuron is determined by the stronger input of the two sources, and the mismatch between these inputs is used to update the synaptic weights of the CA1-dSub pathway. By reproducing the neural dynamics of dSub neurons reported by Poulter et al. 2021, our model suggests a specific role for subiculum in spatial memory: the updating of associative memory (here, between spatial locations and environmental boundaries). This function may also extend to non-spatial settings and provide a framework for integrating the subiculum with the canonical model of hippocampal function.

## 2 Methods

To mimic the task used in the discovery of the vector traces (Poulter et al. 2021), we used the RatInABox toolbox (George et al. 2024) to simulate a virtual rat randomly exploring a 1 m by 1 m square environment with inserted and subsequently removed cues, implementing three trial types: pre-cue, cue, and post-cue trials (the latter manifesting the trace). The virtual rat’s rotational velocity and linear speed were sampled from two different Ornstein Uhlenbeck processes at 100Hz (*dt* = 10ms), and the position, *X*, was updated accordingly (for further details, see George et al. (2024)). An overview of the model structure is shown in Figure 1. The firing rates of dSub neurons are updated based on competing inputs from pSub and CA1. Within this architecture, the main computational function of the CA1-dSub pathway and pSub-dSub interactions is to detect a mismatch between ongoing experience and spatial memory (see learning rule below).

**Figure 1:**
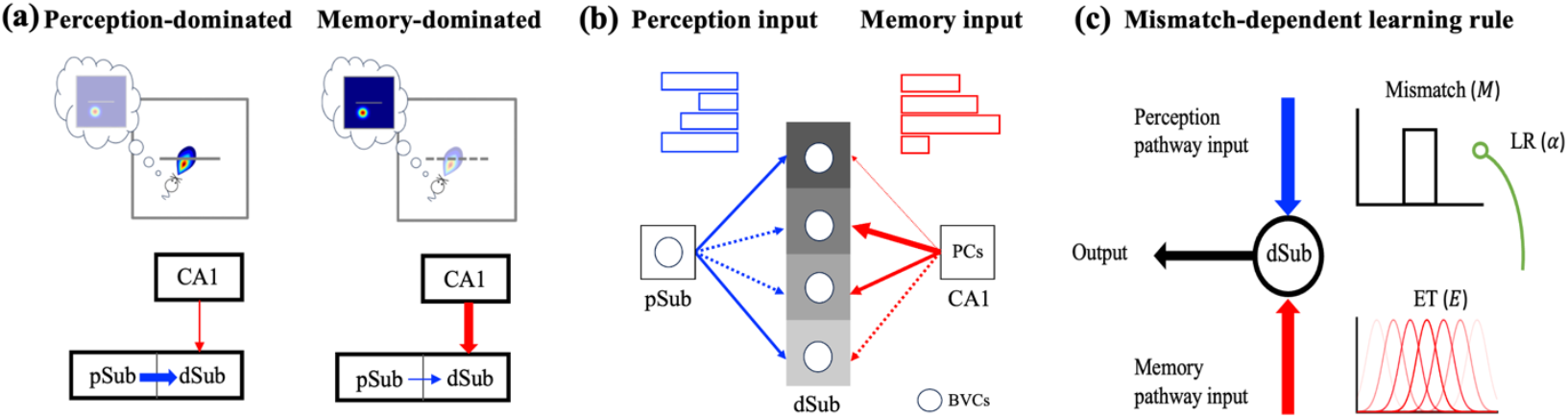
Model schematic. **(a)** Illustration of the network model. The network consists of a perception pathway (pSub-dSub) and a memory pathway (CA1-dSub). Boundary vector cells (BVCs) in the pSub respond to boundaries (gray lines) at a preferred distance and allocentric direction from the virtual rat. Place cells (PCs) in the CA1 respond to the virtual rat’s position. In the perception-dominated mode (left), when a barrier is present in the center, synaptic transmission from pSub is relatively strong (thick blue arrow), and the association between the barrier and the virtual rat’s position is encoded by updating the memory pathway. In the memory-dominated mode, after the barrier is removed, synaptic transmission from region CA1 is relatively strong (thick red arrow), facilitating effective retrieval of previously encoded associative memory. **(b)** Each dSub neuron only receives the stronger signal from either pSub (perception input, blue bars) or CA1 (memory input, red bars). Solid arrows indicate active synapses transmitting information, and dashed arrows represent inactive synapses. The pSub-dSub weights are fixed, while the CA1–dSub synaptic weights are updated at different rates, determined by the learning rates of the postsynaptic dSub neurons, as indicated by distinct colors. Arrow widths represent synaptic strength. (**c)** The learning rule. Coactive neurons generate mismatch signals, computed by subtracting the memory input from the perception input. These mismatch signals are modulated by the learning rates (LRs) of the dSub neurons and are then combined with the eligibility traces (ETs) generated by the CA1 neurons to drive CA1–dSub synaptic plasticity.

### 2.1 The boundary vector cell model

pSub consists of *N*_pSub_ neurons that provide the perception input. The neurons are modeled as non-trace cells (NTCs) that only respond to environmental walls and cues that are present. NTCs map well onto classical BVCs. Each pSub neuron is a simplified model of a BVC (George et al. 2024), with a Gaussian response to the boundaries. The contribution to the firing rates of a pSub neuron *k* (tuning distance *d*_*k*_ and angle *ϕ*_*k*_) from a segment of boundary at distance *r* in allocentric direction *θ*, subtending an angle *δθ* at the virtual rat, is given by:

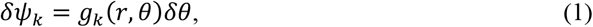

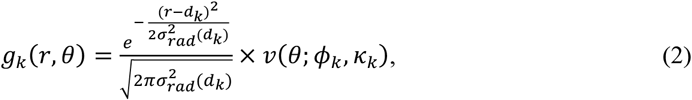

where *v*(*θ*; *ϕ*_*k*_, *κ*_*k*_) is the radial von Mises distribution (a generalization of a Gaussian for periodic variables):

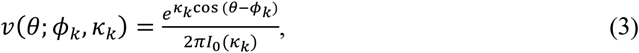

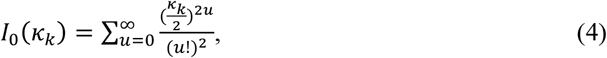

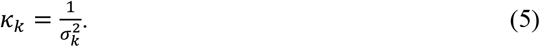

The angular width *σ*_*k*_ follows a uniform distribution, *σ*_*k*_ *∼* 𝒰[10°, 30°]. The radial tuning width *σ*_*rad*_(*d*_*k*_) increases linearly with the tuning distance, 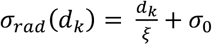, where *ξ* and *σ*_0_ are constants. At time *t*, when the virtual rat is at the location *X*(*t*), the contribution of all boundaries to the firing of the pSub neuron *k* is determined by integrating Eq. (1) over *θ*:

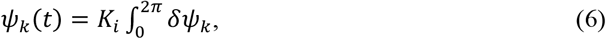

where 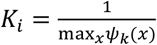 is a normalization constant calculated empirically at initialization such that each BVC has a maximum firing rate of 1.

### 2.2 The place cell model

CA1 consists of *N*_CA1_ PCs that can be thought of as having been initially recruited by grid cell and or some process internal to the hippocampus proper, BVC inputs during first exposure to the environment (Evans et al. 2016; Hartley et al. 2000). This phase is not modelled and PCs are simply given to our model. The firing rate of the CA1 neuron *j*, as a Gaussian tuned response to the virtual rat’s current position, is defined as

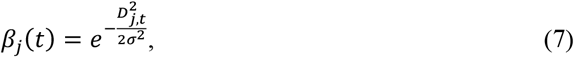

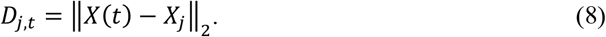

Here, ‖·‖_2_ represents the Euclidean norm of a vector. *X*_*j*_ = (*x*_*j*_, *y*_*j*_) represents the center of the place field of the CA1 neuron *j*. The set of locations {*X*_*j*_}, sampled uniformly at random from the environment, remains constant across the three trials. The width of the place field, *σ*, is constant.

### 2.3 The network

At time *t*, dSub neuron *i* receives memory input currents 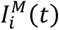 from CA1 neurons and the perception input currents 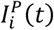 from pSub neurons.

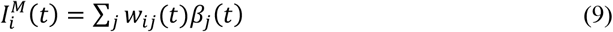

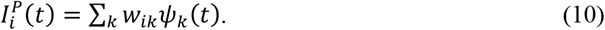

CA1 neuron *j* is connected to dSub neuron *i* through synaptic weights *w*_*ij*_(*t*). The weights of this memory pathway are updated using a mismatch-dependent plasticity rule (see Section 2.4). *k* is the index that labels pSub neurons which are connected to dSub neuron *i*, through synaptic weights *w*_*ik*_. The weights of this perception pathway are fixed, defined as:

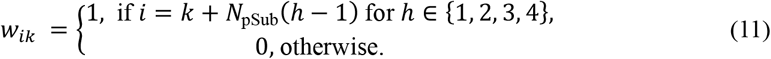

The total number of dSub units is *N*_dSub_ = 4*N*_pSub_. The firing rate *f*_*i*_(*t*) of the dSub unit *i* depends on the larger of the memory and perception input currents, and is computed through a simple sigmoid transfer function.

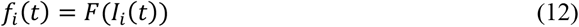

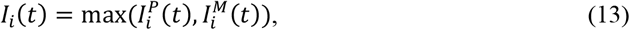

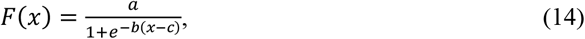

where *F* is a sigmoid function with parameters *a, b* and *c* . Such a max-operation could be implemented in neural terms with lateral inhibition in an intermediate layer, or shunting inhibition (from the stronger input onto the axon conveying the weaker input).

### 2.4 Mismatch-dependent learning rule

Updating of the memory pathway proceeds as follows. We employ a mismatch-dependent learning rule based on the three-factor learning rule (Magee and Grienberger 2020), consisting of the mismatch signal (*M*), eligibility trace (*E*), and learning rate (*α*). At time *t*, the current location *X*(*t*) along the simulated trajectory is determined, and the perception and memory input currents are calculated according to Eq. (9) and Eq. (10), given the current location. Then the activity of all dSub neurons is determined by their sigmoid transfer function (Eq. (12)). The weights are then updated according to a mismatch-dependent learning rule. The mismatch signal is defined as the difference between the perception and memory input currents, given by

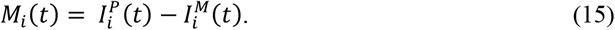

Before learning, the weights are initialized as *w*_*ij*_(0). During learning, the synaptic weights *w*_*ij*_(*t*) are updated at intervals of *τ*,

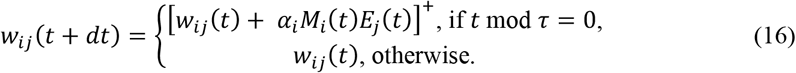

After each updating, the weights are further clipped to between 0 and 1 by the threshold-linear function [·]^+^ . dSub neuron *i* has learning rate, 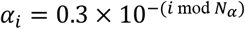, where *N*_*α*_ denotes the number of distinct learning rate values. *E*_*j*_(*t*) is the eligibility trace, an internal signal local to the synapse and independent of dSub output activities, which could for instance be generated by CA1. We first simulate the simplest possible model, where the firing rate of CA1 neuron *j* simultaneously determines both the memory input currents (Eq. (9)) and the eligibility trace *E*_*j*_, defined as

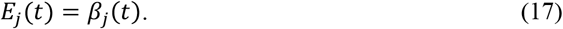

The next section outlines how a more complex eligibility trace affects the model.

### 2.5 Model with cue-dependent PC modulation

For this model variant (Figures 4-6), we assumed that the novelty information from an inserted cue affects CA1 differently along the proximodistal axis. Therefore, the CA1 module is divided into two parts: dCA1 and pCA1, each containing *N*_CA1_ PCs. The firing rates of dCA1 neurons, 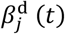, influences learning rule (Eq. (16)) by determining the value of eligible trace:

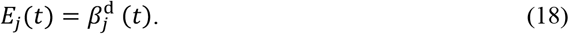

pCA1 neuron *j* is connected to the dSub neuron *i* through *w*_*ij*_. The firing rates of pCA1 neurons, 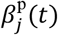, provide the memory input currents,

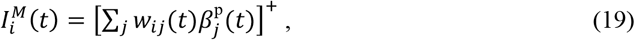

which can also influence the learning rule (Eq. (16)) by determine the value of the mismatch signal (Eq. (15)). The tuning functions of dCA1 and pCA1 place cells are only affected by the inserted cue, reminiscent of experimental findings (Bourboulou et al. 2019; Ormond and O’Keefe 2022; Sato et al. 2020; Turi et al. 2019; Vandrey et al. 2021). During the pre-cue and post-cue trials, the tuning functions are assumed to be unaffected and remain the same as mentioned above:

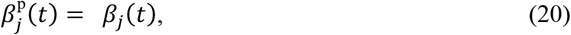

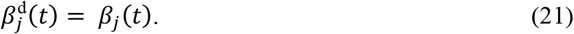

Next, we discuss the tuning functions of dCA1 and pCA1 during the cue trial.

#### 2.5.1 dCA1 place field modulation

We assume that when the virtual rat perceives the presence of the cue at time *t*, the place field centers of the dCA1 neurons would shift by a vector, *V*(*t*). We define *V*(*t*) as the shortest vector pointing from the current position of the virtual rat (*X*(*t*)) to the cue, and the firing rate of the dCA1 neuron *j* is given by

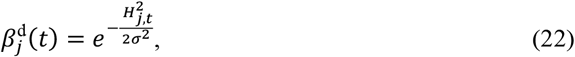

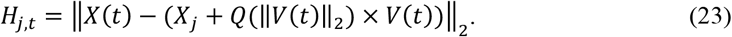

Here, *Q* is a nonlinear function with parameters *m*_1_ and *m*_2_, given by

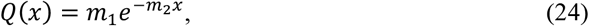

which shows that the shifting of dCA1 place centers is influenced by the distance between the virtual rat and the cue. *Q* is a key ingredient to explain richer dynamics, such as the observed changes of place fields with local effects (Figure 4), which is used to model the shift of the trace field compared to the cue field.

#### 2.5.2 pCA1 place field modulation

The inserted cue influences the pCA1 activity by scaling the amplitude of the tuning functions. The firing rate of the pCA1 neuron *j* is boosted when the cue is within the field of view and attenuated otherwise, defined as:

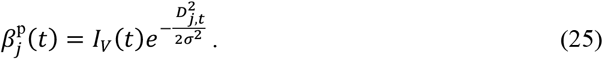

Here, *I*_*V*_ denotes the visual signal from the inserted cue. We set *I*_*V*_ = 100 when the cue is within both the viewing angle range (90°) and distance range (1m), and *I*_*V*_ = 0.001 otherwise.

### 2.6 Simulation details

We began by studying the model with the basic components: a network of dSub neurons receiving input from pSub BVCs and CA1 PCs (Sections 2.1–2.3), together with a synaptic learning rule between dSub and CA1 (Section 2.4). In each trial, the virtual rat explored the environment for 20 minutes. To analyze the effect of cue size on the trace field, we simulated the virtual rat exploring environments containing barriers of varying lengths (Figure 3).

**Figure 2:**
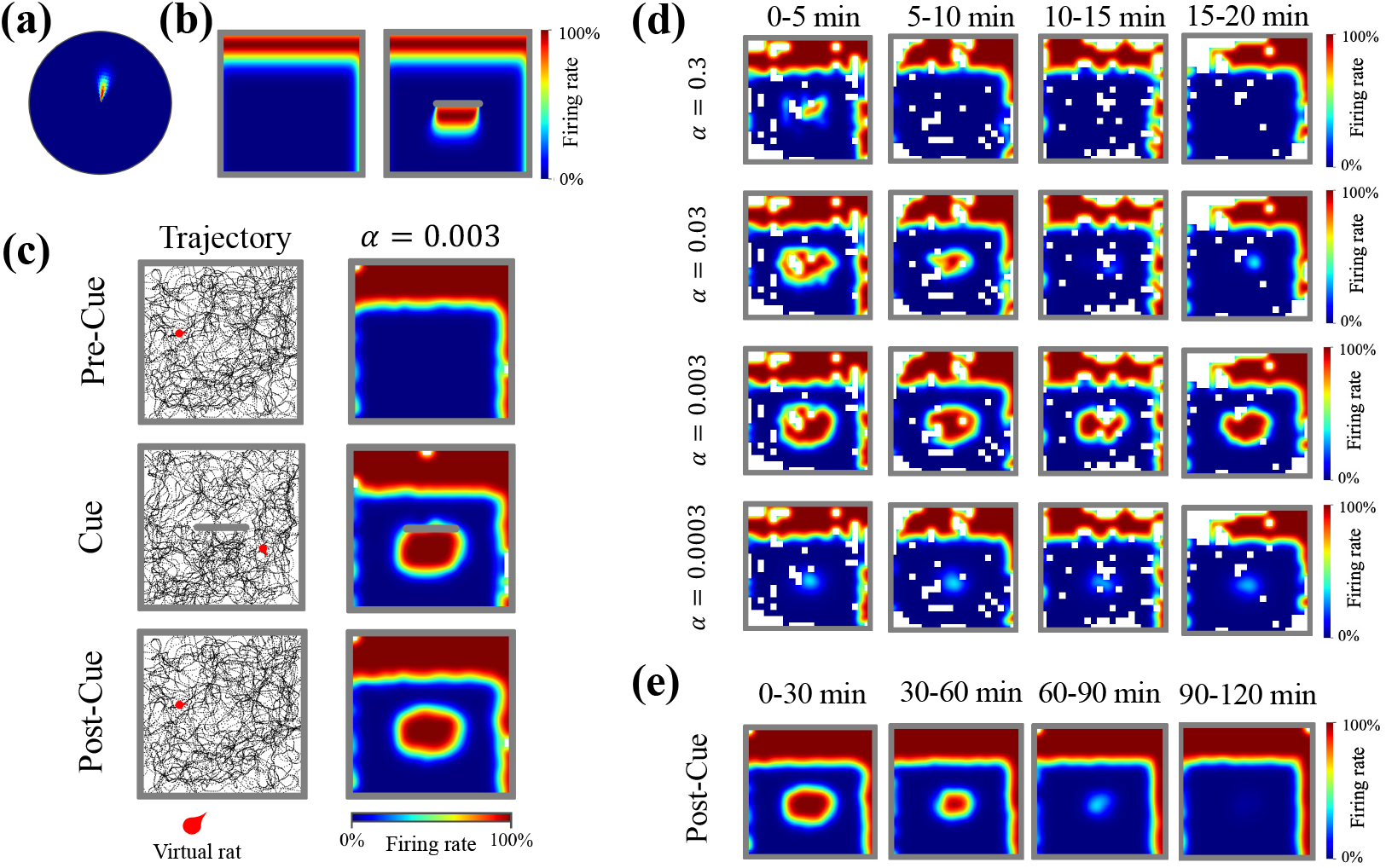
Neural activity of subiculum neurons. (a) A vector plot of the example pSub neuron, which shows the mean firing rate as a function of the virtual rat’s displacement from the boundary. In this example, the neuron fires maximally when a boundary is located 8.90 cm (tuning distance) away at an angle of 81.97° (tuning angle) relative to the virtual rat. Angles are measured clockwise, east = 0°. (b) Rate maps of the example pSub neuron in environments with (left) and without (right) an inserted cue. (c) Example exploration trajectories and rate maps of the dSub neuron with intermediate learning rate (*α*) across three trials. The dSub neuron receives perception input from the example pSub neuron (a, b). (d) Dynamics of trace fields of example dSub neurons with different learning rates (*α*) in the post-cue trial. *α* affects both the size and the decay rate of the trace fields. White bins represent locations that were not visited by the virtual rat. (e) Dynamics of trace fields of dSub neurons (*α* = 0.003) during the post-cue trial over two hours.

**Figure 3:**
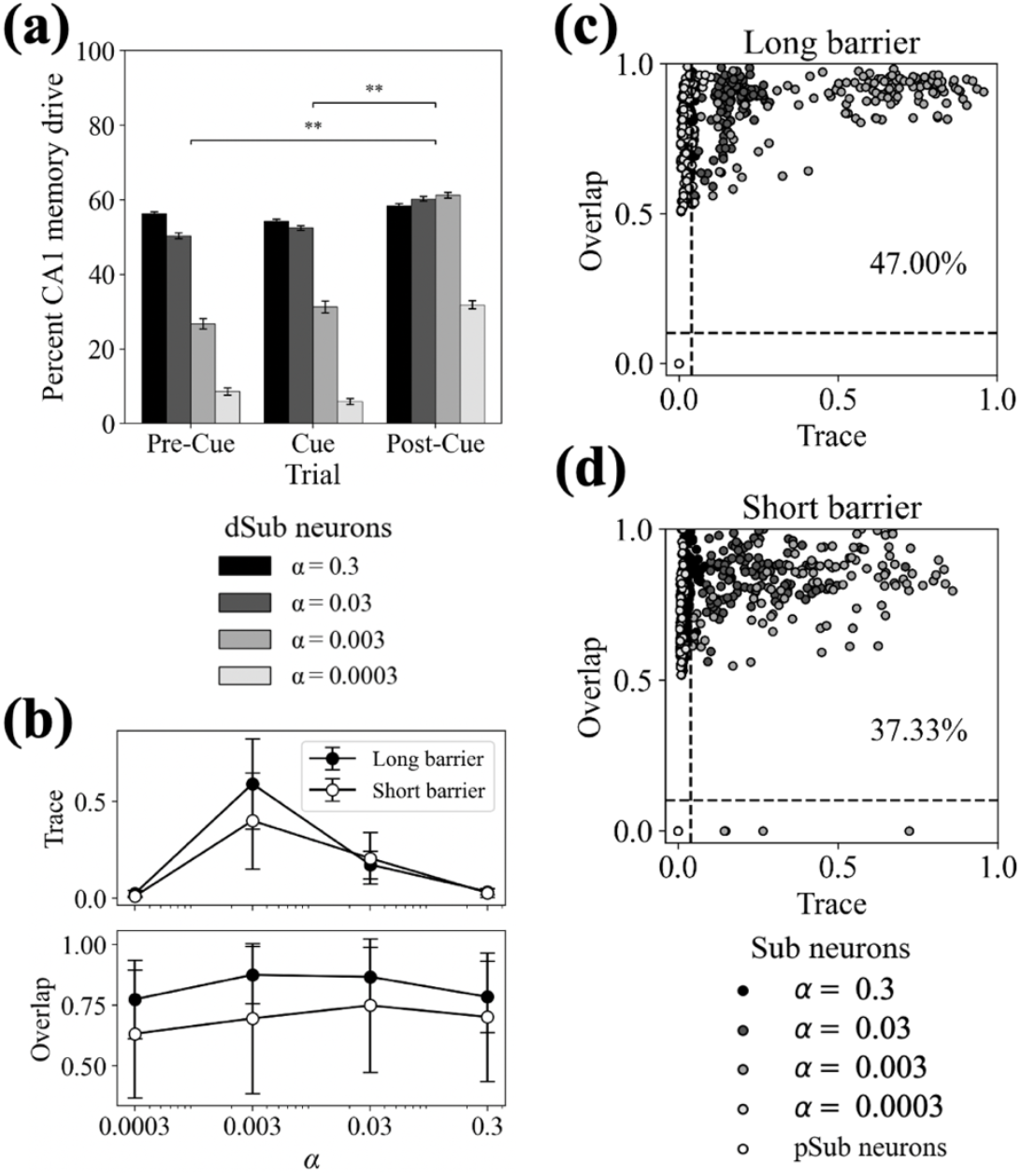
Characterization of the subiculum neurons. (a) Percentage of time during which dSub neurons receive CA1 memory inputs in the simulation. No effect of learning rate (*α*) was found when analyzing the dSub neurons between pre-cue and cue trials, 𝒳^2^(3) = 0.928, *p* = 0.8186 > 0.05. Significant effect of *α* was found when comparing dSub neurons between pre-cue and post-cue trials, 𝒳^2^(3) = 14.538, *p* = 0.0023 < 0.01, as well as between cue and post-cue trials, 𝒳^2^(3) = 15.844, *p* = 0.0012 < 0.01. The data are shown as mean ± standard error of the mean. ** *p* < 0.01, *** *p* < 0.001. (b) Trace and overlap scores of dSub neurons for different learning rates (*α, n* = 120). The data are shown as mean ± standard error of the mean. (c-d) Scatter plots of trace and overlap scores for all subiculum neurons. Vector trace cells (VTCs) were defined based on combined above-threshold trace and overlap scores. (c) 47.00% of subiculum neurons were classified as VTCs with a long barrier (30cm) inserted during the Cue trial. (d) 37.33% were classified as VTCs with a short barrier (10cm). Dashed lines indicate threshold values.

Subsequently cue-dependent modulation of CA1 activity was introduced (Section 2.5). We first simulated the model with only pCA1 place field modulation (Section 2.5.1), using the same 20-minute duration for each trial. The output of the dSub neurons under this condition differs from that of the model without cue modulation (see Eq. (13)). The input currents that determine the firing rate of the dSub unit *i, I*_*i*_(*t*), is defined as:

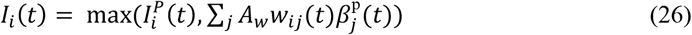

*A*_*w*_ is a constant representing the maximum efficacy of the memory weights, which ensures scale consistency with the perception input currents from pSub.

Finally, we also simulated the model with both pCA1 and dCA1 place field modulation (Sections 2.5.1-2.5.2). In this setting, the virtual rat requires more exploration time to establish associative memory between dSub neurons and CA1 PCs, thereby generating trace fields. This is necessary because the view-dependent memory query boosts the memory pathway, which decreases the total time spent learning. We used 60-minute duration for the pre-cue and cue trial, and 20-minute duration for the post-cue trial. The input currents that determine the firing rate of the dSub unit *i, I*_*i*_(*t*), is defined as:

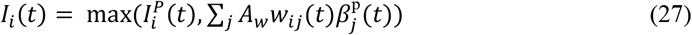

We assume that the pCA1 modulation (a boost in firing rate) captures addition excitation allocated to a cue to facilitate memory readout. Therefore, we compare 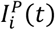 with 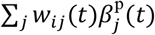 directly, rather than using the memory input currents with a threshold function (Eq. (19)).

### 2.7 Rate maps

Rate maps for all simulated neurons were generated by first dividing the environment into a grid of 5 cm by 5 cm square bins, and then computing the mean firing rate in each bin based only on the firing observed when the virtual rat visited that bin.

### 2.8 Cue fields and trace fields

To evaluate rate maps, we defined cue fields and trace fields according to Poulter et al. (2021). To define cue fields, we first subtracted the pre-cue rate map from the cue-trial rate map. Cue fields were then defined as contiguous regions of the resulting map with a value ≥ 0.75. For trace fields we first subtracted the pre-cue rate map from the post-cue rate map. Trace fields were then defined as contiguous regions of the resulting map with a value ≥ 0.2. For both definitions, if multiple fields were present, only the largest one was used for analysis. Additionally, small trial-to-trial variations in firing rates (e.g., due to movement parameters) sometimes trigger mismatch for wall fields. We assume a threshold (i.e., minimum change) needed to initiate mismatch learning would preclude such effects. To focus our analyses on the cue fields, we excluded bins located within 10 cm of the east or west walls and within 20 cm of the north or south walls. In our simulations, dSub neurons tuning to the north or south were more likely to generate trace fields, as the cue was presented parallel to the north and south walls. Bins that were not visited by the virtual rat during the pre-cue or cue trial were deleted from the analysis.

### 2.9 Trace and overlap scores

The trace score was defined as the average firing rate within the cue field (see Section 2.8) during the post-cue trial, divided by the average firing rate within the cue field during the cue trial. A trace score of 1 indicates that the response based on memory retrieval is as strong as the response in the presence of the cue. The overlap score measures how much the trace field (see Section 2.8) firing overlaps with the cue field firing. If the trace field existed, the overlap score was defined as: 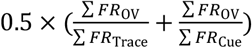 Here, ∑ *FR* represents the total firing rate within a region of the post-cue rate map: ∑ *FR*_Cue_ is the summed firing rate in the cue field, ∑ *FR*_Trace_ is the summed firing in the trace field, and ∑ *FR*_OV_ is the summed rate in the area where the cue and trace fields overlap. If no trace field was detected, the overlap score was set to zero. We adopted the definitions of the two scores from the empirical study (Poulter et al. 2021), with the exception that the firing rates of cue and trace fields were not normalized.

### 2.10 Cue-related tuning distances

We calculated the tuning distance of each dSub neuron in response to the cue. For the cue trial, we first computed the average coordinates of the cue field, weighted by the firing rate in each bin. Since the cue was modeled as a line parallel to the southern and northern walls, the tuning distance in the cue trial was defined as the absolute difference between the resulting y-coordinate and *y*_cue_. For the post-cue trial, tuning distance was computed from the trace field using the same procedure.

## 3 Results

Our CA1–subiculum network model is tested on the behavioral task of ref.(Poulter et al. 2021). Neural activities in the dorsal subiculum were recorded during three trials: before cue insertion (pre-cue trial), during cue insertion (cue trial), and after cue removal (post-cue trial). In our model, the output of the dSub neurons is determined by the competition between perception inputs from pSub BVCs and memory inputs from CA1 PCs. Together with the mismatch-dependent learning rule, the model accounts for multiple characteristics of VTCs (Section 3.1-3.3). We then expanded the model to explore how a dynamic CA1 representation, reflecting the effects of inserted objects and rewards on PCs (Bourboulou et al. 2019; Sato et al. 2020; Turi et al. 2019), manifests in the model (Section 3.4-3.6).

### 3.1 Network model of the CA1-subiculum pathway

The network structure can be seen as complementary to the classic BVC-to-PC model (Hartley et al. 2000) in which PCs are driven by BVC (subiculum to CA1). I.e., we do not suggest that the BVC-PC pathway is not present (see Introduction). During the cue-trial the associative memory of the cue and spatial locations is encoded in the weights between CA1 PCs and subiculum BVCs. After cue removal (post-cue trial), the generation of trace field is driven by the inputs from the PCs through the CA1-subiculum pathway (Figure 1a). Here we considered differences in BVCs along the proximodistal axis of the subiculum, a hitherto unexplored aspect. pSub mainly includes NTCs, while dSub neurons are composed of both VTCs and NTCs (Poulter et al. 2021). We equate pSub BVCs to NTCs in our model (Figure 2a-b). These cells only show cue-related firing in the presence of the cue and we interpreted this firing as a result of upstream perception (suitably transformed to allocentric format (Alexander et al. 2023; Bicanski and Burgess 2018; Byrne, Becker and Burgess 2007)). In our model, dSub neurons also receive perception inputs from pSub BVCs. This aligns with observations that pSub can transmit information to dSub, but not vice versa (Matsumoto, Kitanishi, and Mizuseki 2019). The firing rates of dSub neurons are determined by the higher input value between CA1 PCs and pSub NTCs (Figure 1b). This competitive mechanism enables both NTCs and VTCs in dSub to respond to the present cue, as dSub can continuously receive perception inputs from pSub NTCs. However, after cue removal, only dSub neurons that have learned the association to place (via related PCs) during the cue trial can generate the trace field (see Section 2.8).

The updating of associations depends on the mismatch-dependent learning rule (Figure 1c; Section 2.4). CA1 PC firing serves as the eligibility trace. Thus, it not only triggers the stored memory but also influences current memory updating. Each dSub neuron produces a mismatch signal by calculating the difference between perception inputs and memory inputs. As a result, the CA1-dSub pathway is trained to incrementally reduce the discrepancy between external perception and internal memory with varying learning rates. We can build an intuition for the memory updating process as follows. In the pre-cue trial, both perception and memory inputs for the cue are absent, indicating that no memory updating occurs. Due to the introduction of the cue, the corresponding perception inputs lead to a positive mismatch signal, indicating a memory formation process. After cue removal, perception inputs (for the cue) disappear, however, the memory inputs persist due to the learning that has occurred in the cue trial. This leads to a negative mismatch signal, indicating unlearning of an association. The mismatch signal and eligibility trace are related to current perception inputs from the external world, depending on the virtual rat’s exploratory behavior and the environment setup. The learning rate is specific to each dSub neuron, and we assume it captures individual differences among dSub neurons, specifically between NTCs and VTCs.

### 3.2 Vector trace driven by memory inputs after learning

The basic model (see Sections 2.1–2.4), reproduces the emergence of trace fields and most of the experimental findings on VTCs (Poulter et al. 2021). Our analysis focuses on the output of dSub neurons to determine the factors influencing the trace field. Figure 2 shows examples of activity patterns of dSub neurons with different learning rates obtained from the model. We found that dSub neurons and the connected pSub BVCs (Figure 2a-b) exhibit similar tuning directions and distances to environmental boundaries during pre-cue and cue trials (Figure 2c). In the post-cue trial, the dSub neuron with an intermediate learning rate (*α* = 0.003) exhibited the strongest trace field (Figure 2c).

Trace fields result from a balance between memory updating and retention (Figure 2d). When the learning rate is relatively large, the trace field was initially significant but decayed very rapidly, indicating efficient memory updating. The associative memory is well-established after the cue trial, but it is also quickly erased when the cue is no longer present. Conversely, when the learning rate is relatively small, associative memory is not well-established after the cue trial, resulting in a weak initial trace field. However, the constructed trace field can be maintained for a longer duration, indicating robust memory retention. dSub neurons with an intermediate learning rate exhibited a significant trace field that persisted stably for the first 20 minutes. We also ran a simulation where the virtual rat explored for a sufficiently long duration during the post-cue trial, and found hours-long persistence of trace field (Figure 2e), aligning with experimental observations (Poulter et al. 2018). We suggest that this spread in learning rates allows the system to differentiate between cues with respect to spatial size and temporal persistence, capturing a wide range of possible associations.

### 3.3 The role of dSub neurons in memory retrieval

To examine how the drive to individual neurons changes, we analyzed the source of dSub neuronal activity across the three trial types (Figure 3a). We found that there was no significant difference in the distribution of neuron types between the pre-cue and cue trials. Moreover, boundary-related memory information was consistently represented by neurons with larger learning rate in these two trials. This demonstrates that more certain boundary settings—those consistent with current perception—were mainly represented by neurons with rapid memory updating, thereby reducing the time required for memory formation. However, in the post-cue trial, dSub primarily recruited neurons with relatively smaller learning rate to represent memory information in the post-cue trial. I.e., cue-memory inconsistent with current perception was mainly represented by neurons with slower updating, prolonging memory retention.

In experiments, dSub neurons showed sensitivity to varying cue types, exhibiting different proportions of VTCs and NTCs when exposed to distinct cue types (Poulter et al. 2021). We thus ran simulations in which the virtual rat explored environments containing barriers of different lengths. dSub neurons were classified into VTCs and NTCs based on trace score and overlap score. (Figure 3b; see Section 2.9). The trace score quantified the strength of cue-related firing during the post-cue trial, while the overlap score measured the spatial overlap of cue-related firing between the cue and post-cue trial. For pSub neurons, which were directly modeled as NTCs, both trace and overlap scores were zero. dSub neurons with trace scores ≥ 0.04 and overlap scores ≥ 0.1 were classified as VTCs; otherwise, they were considered NTCs.

We found that larger cue sizes elicited more trace responses (Figure 3c-d). Larger cues occupied a greater area in the environment, increasing the likelihood that the virtual rat explored near the cue, suggesting more candidate PCs can establish connections with dSub neurons. dSub neurons therefore exhibited lower overlap scores in the short-barrier condition (Figure 3b), suggesting that associative memory was less well established. Moreover, we found that more dSub neurons with larger learning rates were classified as VTCs in short barrier condition (Figure 3d), implying that a faster updating rate promotes memory retrieval under limited exploration.

### 3.4 Cue-dependent modulation in CA1

The above model with basic elements already captures the majority of findings on VTCs (Poulter et al. 2021). However, the inserted cue introduces novel information, which significantly impacts exploratory behavior and hippocampal representations (Akiti et al. 2022; Cen et al. 2024; Dong et al. 2021; Shen and Dayan 2024). Since we assume that the trace field is driven by CA1, we introduced cue-dependent modulation of CA1 PC activity to explore its effect on the trace field (see Section 2.5). Both theoretical and experimental studies demonstrated CA1 place fields shifting towards an object/reward location (Bourboulou et al. 2019; Gerstner and Abbott 1996; Grienberger and Magee 2022; Kaufman, Geiller and Losonczy, 2020; Kecket al. 2024; Sato et al. 2020; Sosa et al. 2023; Turi et al., 2019). In the CA1-subiculum neural circuit, dCA1 has been theorized to convey more object-related information, while pCA1 is thought to transmit mainly spatial information (Igarashi et al. 2014; Knierim, Neunuebel, and Deshmukh 2014; Nagelhus et al. 2023; Nakazawa et al. 2016; Vandrey et al. 2021). Thus, we assumed that the inserted cue affected the place fields of dCA1 PCs in our model (Figure 4a; see Section 2.5.1). On the linear track, CA1 place fields near the object/reward will shift, whereas place fields farther away are less affected (Bourboulou et al. 2019; Sato et al. 2020; Turi et al. 2019). By using a negative nonlinear map of the distance between cue and the virtual rat’s position (see Section 2.5.1), we implemented the localized shift (Figure 4c) closely resembling experimental data (Figure 4b; Turi et al. 2019).

**Figure 4:**
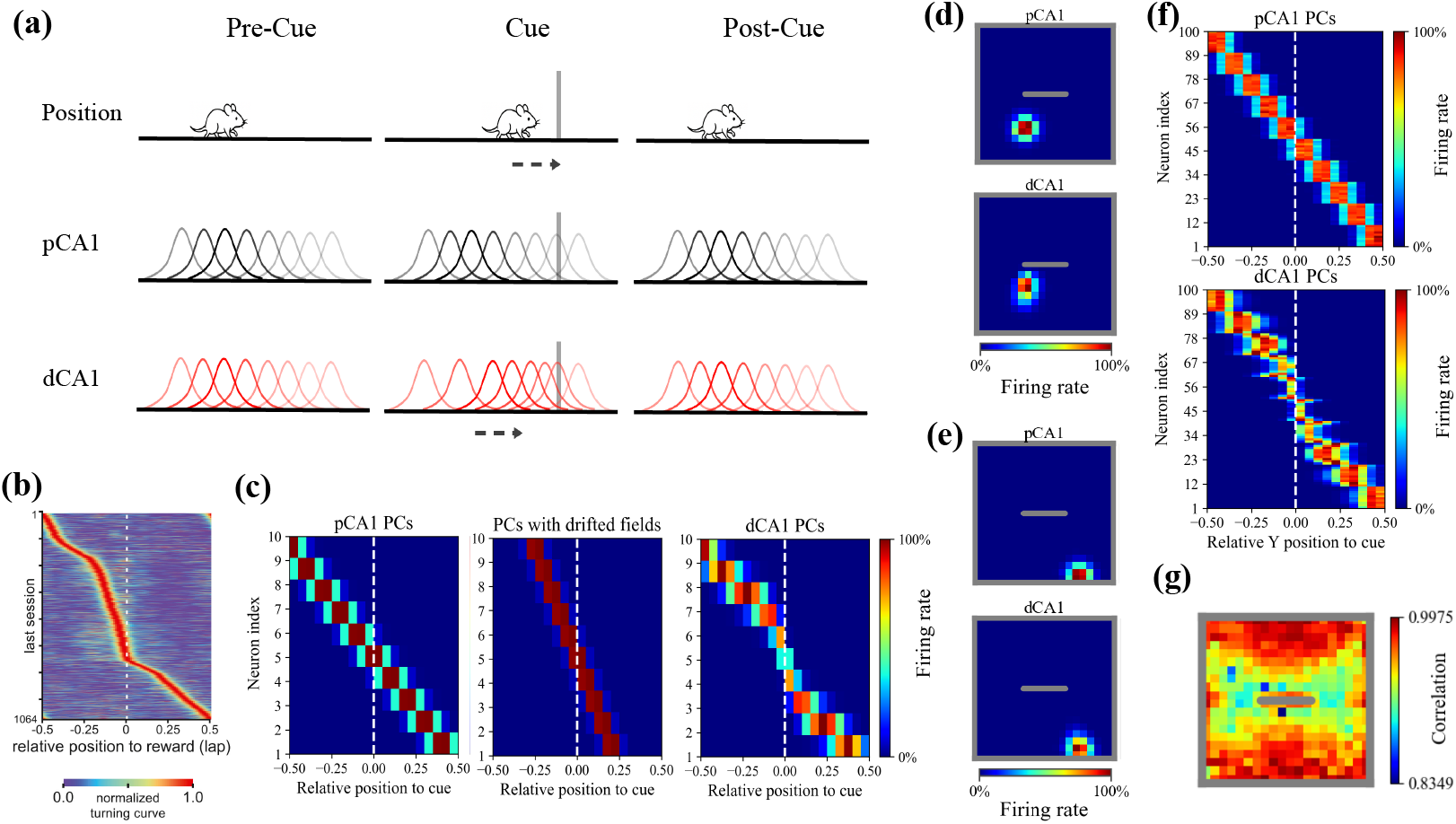
CA1 place field modulation in response to a cue/reward. (a) Schematics illustrating features of CA1 representations on a 1D track. Top: A virtual rat on a linear track. The grey bar represents the inserted cue, and the dashed arrow represents *V*, the shortest vector from the rat’s position to the cue. Middle: Gaussian functions represent pCA1 place cells (PCs) with fixed distributed place fields across three trials. Transparency indicates firing rate values. Bottom: Gaussian functions represent dCA1 PCs with place fields shifting towards the cue according to *V* (see Section 2). (b) Experimental evidence for the local shift of CA1 place field. On a 1D treadmill belt, PCs became more concentrated near the reward zone after reward learning (see slope close to dashed line), whereas PCs farther away were less affected. (b) Adapted from Turi et al. (2019) with permission. Similar effects have been shown for objects (Bourboulou et al. 2019; Sato et al. 2020). (c) Example of simulated place field distributions in a 1D environment during the cue trial. Left: pCA1 place fields are uniformly distributed. Middle: Using only *V*, all place fields shift toward the cue leaving the edges uncovered. Right: dCA1 place fields exhibit local shift toward the cue, by using the combination of a nonlinear function *Q* and *V, Q* × *V* (see Section 2). (d,e) Example single CA1 activity in a 2D environment. The example dCA1 and pCA1 PCs share the same place field in the environments without the cue. (d) Rate maps of CA1 PCs located near the cue. (e) Rate maps of CA1 PCs located far from the cue. (f) Example of simulated CA1 place field distributions in 2D environment. The coordinates of place fields were projected onto 1D, using the Y-axis. (g) Correlation map showing the similarity of time-averaged CA1 population activity between simulations with and without the cue. Correlations are higher in the periphery. Compare to Figure 4 in Nagelhus et al. 2023.

We then augmented our model with proximodistal differences of CA1. Figure 4d-f illustrates the features of the CA1 region in the augmented model. In the 2D environment, pCA1 place fields remained consistent across the three trials. The dCA1 place fields were consistent with pCA1 place fields during both the pre-cue and post-cue trials. However, in the cue trial, dCA1 place fields shifted locally toward the inserted barrier (Figure 4d-e), consistent with experimental findings that dCA1 place fields near objects are more likely to remap (Vandrey, Duncan, and Ainge 2021), and that the average field size decreases in presence of objects (Burke et al., 2011). At the population level, dCA1 place fields demonstrated localized shifting in the 2D environment reminiscent of the 1D environment (Figure 4f). We also computed the pairwise Pearson correlation between stacks of rate maps of all CA1 PCs, including both pCA1 and dCA1 cells, from the pre-cue and cue trials. This resulted in a correlation map (Figure 4g), where each spatial bin had a correlation value, indicating that PCs responded to the cue based on the virtual rat’s distance from the cue. Similar results have been observed in the CA1 region (Nagelhus et al. 2023). Figure 5a illustrates the structure of the augmented model. In this model, pCA1 PCs project to dSub neurons and provide memory inputs, consistent with the known anatomical connectivity of the pCA1-dSub pathway (Cembrowski et al. 2018; Knierim, Neunuebel, and Deshmukh 2014; Matsumoto et al. 2019; Sun et al. 2018). The firing of dCA1 PCs serves as an eligibility trace in the mismatch-based learning rule (see Section 2).

**Figure 5:**
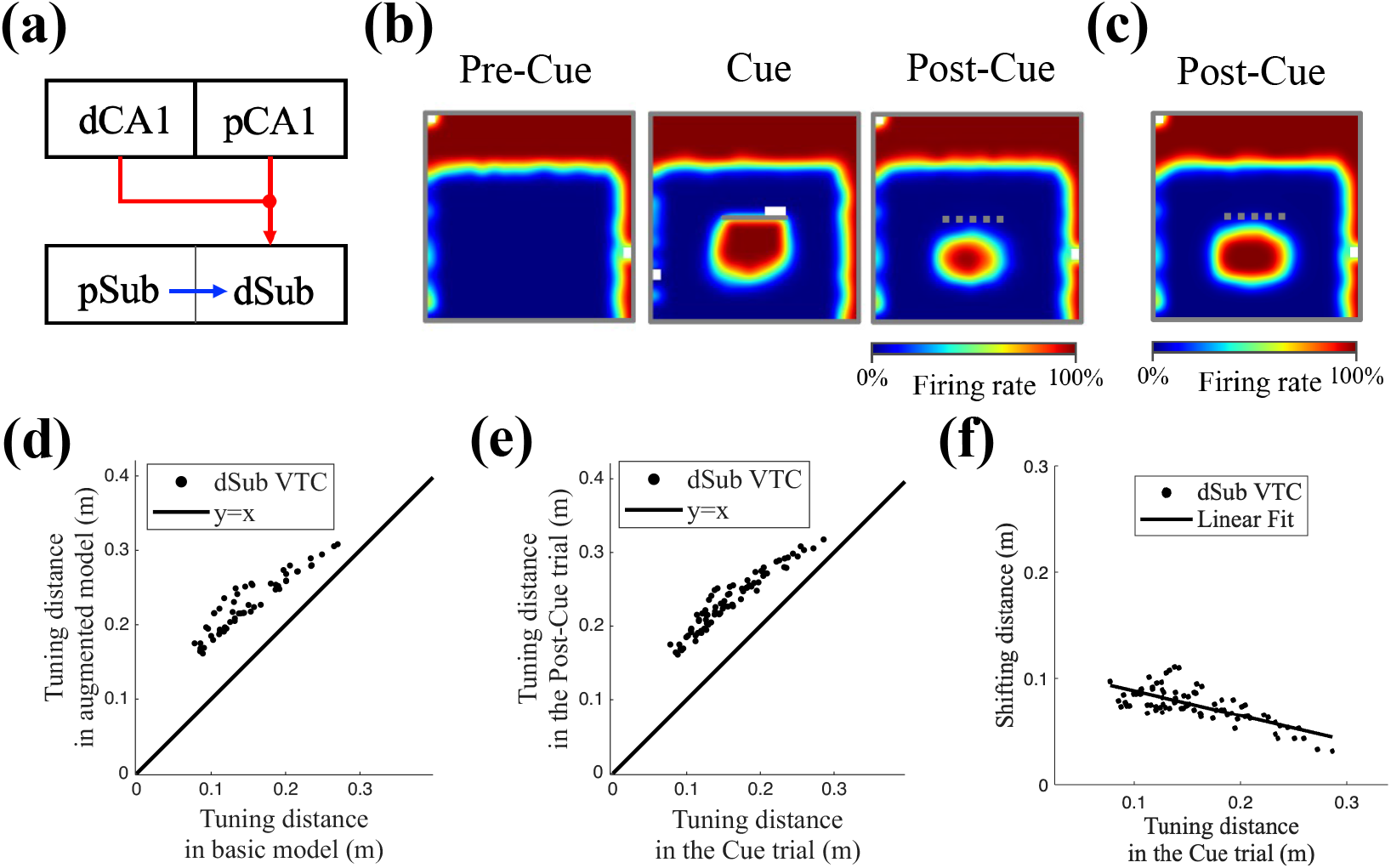
dCA1 place field modulation causes shifts in trace field locations in the post-cue trial. (a) Illustration of the network model incorporating proximodistal differences in CA1. The same network structure as the model shown in Fig. 1a (see Section 2.1-2.4), except that pCA1 place cells (PCs) provide memory inputs, while dCA1 PCs activity represents the eligibility trace (*E*) that modulates synaptic plasticity of pCA1-dSub pathway (see Section 2.5.1). (b) Example rate maps of the dSub neuron with intermediate learning rates (*α* = 0.003) across three trials. (c) Example rate map of the dSub neuron without synaptic plasticity in the post-cue trial. The example neuron was assigned *α* = 0.003 during the pre-cue and cue trials, but *α* was set to 0 in the post-cue trial. (d-e) Scatterplots of cue-related tuning distances for dSub neurons with *α* = 0.003 (n = 82). The definition of cue-related tuning distances is shown in Section 2.10. (d) Comparison of tuning distance in the post-cue trial produced by the basic model (Fig. 1a) and the augmented model(a). (e) Comparison of tuning distance between the cue and post-cue trial. (f) The relationship between shifting and tuning distance during the cue trial. The amount of shift was calculated as the difference in tuning distance between the cue and post-cue trials. Linear regression fit: 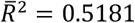.

### 3.5 pCA1 cue-dependent modulation and the effect on dSub neurons

Following validation (Figure 4), we incorporated the proposed cue-dependent CA1 modulation into the proximodistal CA1-subiculum model (Figure 5a) and analyzed the effect of the dCA1 translational modulation (Figure 4d-f, see Section 2.5.1). Consistent with previous results, dSub neurons with intermediate learning rates (*α* = 0.003) generated the strongest trace fields. Importantly, trace fields in the post-cue trial shifted away from the cue relative to the cue-trial (Figure 5b), aligning with experimental findings showing that VTCs exhibited longer tuning distances to the cue in the post-cue trial compared to the cue trial (Poulter et al. 2021). When the virtual rat explored near the inserted cue, all dCA1 PCs shifted their place fields towards the cue (see Section 2). Therefore, dCA1 PCs that were initially farther from the cue increased their firing rates, enhancing the likelihood of forming memory connections with the corresponding dSub neurons. Additionally, we ran a simulation with zero learning rate in the post-cue trial, prohibiting the memory erasure in the absence of cue (despite mismatch). We found that the previously reproduced shifted trace fields were still present (Figure 5c), indicating that the trace field shift was a result of learning under dCA1 modulation in the cue trial.

This effect can be interpreted as predictive in nature, allowing the animal to anticipate the presence of the cue from locations farther away than its actual position during perception, when the cue is recalled from memory. To further assess the predictive character, we focused on dSub neurons with intermediate learning rate (*α* = 0.003), given that this group of neurons accounted for the largest proportion of VTCs. The distance a trace field shifted was defined as the difference in cue-related tuning distances between the cue and post-cue trial (see Section 2.10). To ensure that we restricted the analysis to cue-related firing, we excluded shifts ≥ 0.25 which can relate to wall-responses, due to minor variation in firing rate across the three trial types. Similarly, we excluded cells that exhibited inconsistent cue-related tuning directions between the cue and post-cue trials. We first computed the cue-related tuning distances in the post-cue trial as produced by the basic model (Figure 1a) and the augmented model (Figure 5a). We found that dCA1 modulation induced a significant shift of the trace field (Figure 5d). As shown in Figure 5e, VTCs exhibited larger tuning distances in the post-cue trial relative to the cue trial. We then used a linear regression to model the relationship between the shifting distance and tuning distance in the cue trial (Figure 5f). We found VTCs with shorter tuning distances in the presence of the cue showed greater shift, suggesting stronger predictive capability during memory retrieval.

### 3.6 dCA1 cue-dependent modulation and the effect on dSub neurons

Over and above the effect of CA1 memory inputs, we propose that dSub neurons also perform behavior-related predictions when receiving the perceptual information about the cue. This was prompted by the suggestion (personal communication, Colin Lever), that VTC firing fields, split by HD, exhibited what can be interpreted as direction-modulated anticipatory firing. That is, the cue field locations appear shifted relative to each other in the direction opposite of current HD during the cue trial. I.e. when an animal moved eastward, the cue field appeared further westward. Compared to the trace field shift in the post-cue trial (Figure 5b), this directional shift implies a predictive map that can guide the animal’s movement. HD information has been found in the subicular complex (Simonnet and Fricker 2018; Simonnet et al. 2017).

To model this effect, we incorporate both dCA1 and pCA1 modulation (Figure 6a). We assumed the amplitude of pCA1 firing increases, providing stronger memory inputs, when the cue is within the virtual rat’s field of view (*I*_*V*_ > 1), and the amplitude decreases when the cue is not in view (*I*_*V*_ < 1). This is inspired by the observation that some CA1 PCs respond maximally to the goal in front of the animal (Ormond and O’Keefe 2022; Sarel et al. 2017). The introduction of this pCA1 modulation leads to a HD-related shift of the cue field (Figure 6b), and we interpreted this pCA1 modulation as a memory query (e.g., “was this thing I am looking at here before?”), triggering memory readout of dSub neurons upon receiving the visual signal of the present cue. In our simulation, HD was binned according to 4 quadrants/90-degree cones (facing North, East, South, West). HD was related to the value of *I*_*V*_. For example, a dSub neuron connected to a pSub NTC with a tuning direction of North. When the HD was East, *I*_*V*_ > 1 if the virtual rat was positioned Southwest of the cue. The dSub neuron thus received strong pCA1 memory inputs but weak pSub perception inputs. In contrast, *I*_*V*_ < 1 if the virtual rat was Southeast of the cue, then both perception and memory inputs were weak. The combined influence of the virtual rat’s HD and the cue’s spatial location caused the trace field to shift Westward. Additionally, *I*_*V*_ also influenced synaptic plasticity in the pCA1–dSub pathway by modulating the mismatch signal (see Section 2). When *I*_*V*_ > 1, the mismatch signal was reduced due to enhanced memory input, thereby hindering the rate of establishing associative memory. Therefore, we prolonged the simulation time for the pre-cue and cue trial to generate the trace field in this model-variant (see Section 2.6). Note that this HD-related effect manifests in the cue trial. Consistent with the experimental observation (Poulter et al. 2021) and our previous results (Figure 5b), we still observed the shift of the trace field (cue vs post-cue trial) in the post-cue trial (Figure 6c).

**Figure 6:**
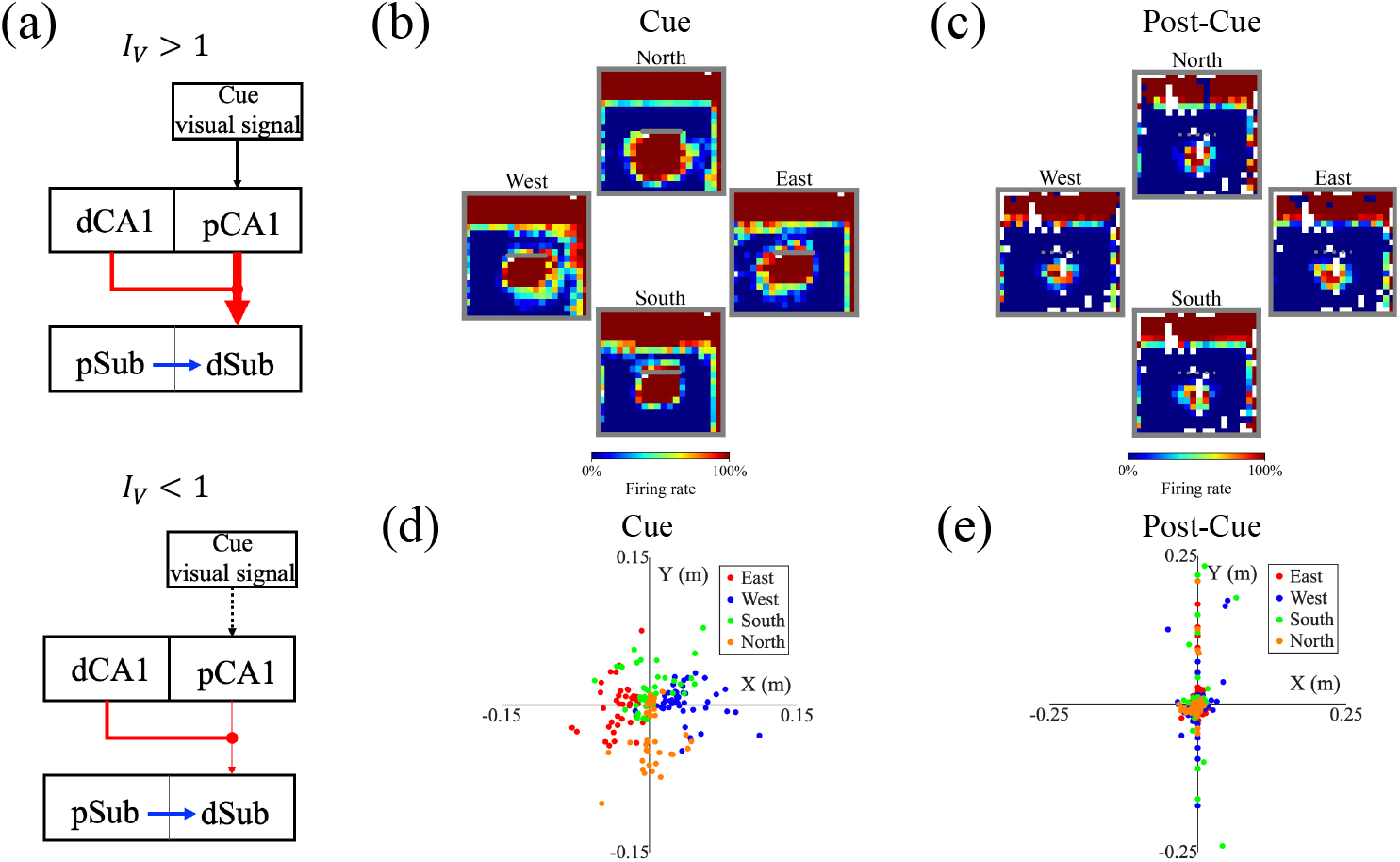
Head direction-related anticipatory shifts of trace fields in the cue trial. (a) Illustration of the network model incorporating both dCA1 and pCA1 place field modulation. Top: When the cue is within the virtual rat’s field of view, memory inputs are strengthened through enhanced firing of pCA1 PCs. Bottom: When the cue is outside the field of view, memory inputs are suppressed through reduced firing of pCA1 PCs (see Section 2). Other settings are consistent with the model shown in Figure 5a. (b-c) Example rate maps of dSub neurons with a learning rate of *α* = 0.003, classified by head direction (HD). HD was categorized into four groups based on angle ranges: eastward (0°-45° and 315°-360°), northward (45°-135°), westward (135°-225°), and southward (225°-315°). Angles are measured clockwise, with 0° indicating the eastward direction. (b) Cue trial. (c) Post-cue trial. (d, e) Coordinate representation of shifting vectors. (d) Cue trial: Each shifting vector points from the average coordinates of the cue field to the average coordinates of the HD-grouped cue field. (e) Same as (d) for the post-cue trial. Color indicates HD: red (n = 41), blue (n = 44), green (n = 45), and orange (n = 39).

To characterize this behavior-related predictive capability, we computed the shifting vector that was defined as a 2D vector pointing from the average coordinates of the cue field to the average coordinates of the HD-grouped cue field (Figure 6d). Consistent with above analysis (Figure 5d-f; see Section 3.6), we focused on dSub neurons with *α* = 0.003. The shifting distances and tuning directions were computed based on rate maps grouped by the HD. The distribution of shifting vectors in the cue trial indicated that dSub could predict the presence of a cue in the upcoming direction. We also computed the post-cue shifting vectors based on the trace fields and found no modulation from HD (Figure 6e).

## 4 Discussion

We have presented what is – to the best of our knowledge – the first neural model of proximo-distal within-subiculum and CA1 interactions, and suggest that these underly the updating of associative memory between boundaries and locations. We demonstrated that the proximodistal CA1-subiculum network and mismatch learning rule is sufficient to account for the key features of VTCs in the subiculum (Poulter et al. 2021), including the distribution of VTCs along the proximodistal axis; the hours-long persistence of trace field; varying proportions of VTCs and NTCs depending on cue type; the shift of VTC firing fields away from the original cue-elicited location; and variations of firing field location as a function of HD in the presence of the cue (compatible with unpublished data, Colin Lever).

### 4.1 Predictions

The model shows a dedicated role for proximal and distal subiculum (and related framing of proximal and distal CA1) beyond subiculum’s known role in merely conveying output of the hippocampus proper. Notably, within its mechanisms, the model renders consistent the observed effects in CA1 and subiculum. It predicts the potential neural substrates, along with their interactions and functions, involved in memory retrieval. (1) The firing of subiculum BVCs reflects both perception and mnemonic functions, which may map onto cellular differences along the proximodistal axis (Kim and Spruston 2012; Matsumoto et al. 2019; Poulter et al. 2021). (2) Blocking CA1-Sub synaptic plasticity should prevent the formation of the vector trace while potentially leaving BVC activity consistent with the current environmental configuration intact. (3) A range of learning rates is suggested to be functionally beneficial and should be observed in the CA1-subiculum circuit, approximating biologically plausible differences in the magnitude of synaptic potentiation (Strauch et al. 2025); for example, through the distribution of neuromodulatory influence of synaptic modifications (Gu 2002; Magee and Grienberger 2020; Nadim and Bucher 2014). (4) Exploratory movement, which plays a role in updating cognitive maps (Brunec et al. 2023; Manns and Eichenbaum 2009; Redish 1999; Shamash et al. 2023), can affect the recruitment of dSub neurons in memory retrieval. When time is limited (e.g., the animal has fewer opportunities to interact with a short barrier compared to a longer one), neurons with larger learning rate should be recruited. (5) When there is a long pause between the cue trial and the post-cue trial (days), the trace field should still be detected in the post-cue trial, as no mismatch signal occurs during that pause. (6) The inserted cue indirectly modulates dSub activity by altering CA1 activity. CA1 place field shift should be directly correlated with trace field shift. (7) Apart from reward (Grienberger and Magee 2022; Sosa et al. 2023; Turi et al. 2019), dCA1 PCs should also show localized shifts toward other cue types (e.g., barriers, objects, etc.), but the features of the shift may differ across cue types (Bourboulou et al. 2019; Burke et al. 2011; Sato et al. 2020). (8) Short distances are systematically overestimated during recall (Lederman et al. 1987). VTCs with shorter tuning distances in the presence of the cue should exhibit more pronounced trace field shifts. (9) The HD-related receptive field shift during the cue trial is driven by visual input and therefore should not be present in darkness.

### 4.2 Related work

In our model, vector traces in dSub emerge from integrating perceptual and mnemonic information related to boundaries. I.e., information from two sources is combined, a common property in neural systems (Stokes 2015; Yang and Naya 2020). The mismatch between the two sources served as an instructive error signal to facilitating supervised synaptic plasticity. Compared with previous models that employed a pure Hebbian learning rule (Hebb, D. O. 2005) to establish associations between BVCs and PCs (Bicanski and Burgess, 2018; Byrne, Becker, and Burgess, 2007), the mismatch signal facilitates the unlearning in the absence of perception input, thereby reproducing the temporal decay of vector traces. It has been hypothesized that a mismatch signal in the hippocampus can also contribute to memory coding and updating (Hasselmo, Bodelón, and Wyble 2002; Hasselmo and Schnell 1994; O’Keefe 1976). A previous encoding-versus-retrieval model has proposed that the theta rhythm may provide distinct phases in which CA1 receives either perception or memory inputs (Hasselmo et al. 2002). This account focused on CA1-CA3 interactions, precluding memory read-out (from CA3) at one phase of theta and facilitating it at the opposite phase. Thus, this mechanism operated on the indices (place cells), rather than on the indexed content (boundary presence). Interestingly, in the subiculum, distal neurons fire at an earlier theta phase than proximal neurons (Poulter et al. 2021). The theta rhythm might thus be a candidate component in a more detailed neural implementation of our model (Marr 1971). Experimental findings showed that cue insertion elicited a significantly earlier theta phase in dSub BVCs, and this earlier-phase shift was greater in VTCs than in NTCs (Poulter et al. 2021). These phenomena were observed only in the cue field, but not in the wall field, implying that novelty affected the memory updating process.

Numerous studies have demonstrated the influence of novelty on animal exploratory behavior and the formation of cognitive maps (Akiti et al. 2022; Cen et al. 2024; Dong et al. 2021). The theta rhythm in the hippocampal formation has been shown to support novelty detection (Hasselmo, Wyble, and Wallenstein, 1998; Meeter, Murre, and Talamini 2004; Naber, Witter, and Lopes Da Silva 2000; O’Keefe and Nadel 1978), and environmental novelty can elicit a theta phase shift in CA1, but not in the subiculum (Lever et al. 2010). Since CA1 inputs are crucial to our model, this consistency in mechanism between experiments and our model (given the role assigned to CA1) lends credibility to the CA1-subiculum interplay suggested here. We interpret the overexpression of place fields near cues as an enhancement of the spatial resolution for object/reward coding, albeit at the potential cost of reduced spatial resolution far from the cue. Presuming a constant number of recruited PCs per session, this would constitute a trade-off between reliability (more CA1 cells recruited near the cue) and memory precision. Such a trade-off should also naturally limit the extent of overexpression and shift, resulting in a localized shifts of both place fields and trace fields. However, experimental data also indicate that, in the presence of objects, more place cells are present in the dSub, and individual neurons exhibit a higher mean number of fields (Burke et al. 2011). Thus, the underlying mechanisms remain to be explored in more detail in future studies.

The shift of dSub firing fields is reminiscent of psychological phenomena in which distances to viewed objects-in-context are often overestimated during memory retrieval (Mullally, Intraub, and Maguire 2012). This shift could also be interpreted as a predictive function based on memory recall. Neural replay is also a form of memory recall that can predict subsequent behavior, characterized by the sequential reactivation of neuronal populations (Carr, Jadhav and Frank 2011; Foster and Wilson 2006; Ólafsdóttir et al.2015; Wilson and McNaughton 1994). Unlike replay, which typically occurs during rest, the trace field observed in the subiculum reflects a form of real-time, within-trial memory retrieval and dynamic updating of environmental information associated with a cognitive map – i.e., a readout and update of the map during behavior. Previous hippocampal models have described how replay supports planning during spatial navigation (Cazin et al. 2019; Edvardsen, Bicanski and Burgess 2019; Gagne and Dayan 2022). Incorporating the trace field phenomenon may offer insights into how memory retrieval guides navigational behavior online. A predictive map framework has been invoked to explain the emergence of multiple neural representations in the subiculum, including BVCs and corner cells (Bennett et al. 2024). The emergence of corner cells in that framework is intriguing. However, the near-instantaneous within-trial emergence of BVC responses in response to any novel geometry and/or inserted cues in experiments seems less of a match to the slow learning of successor representations. Future work could endeavor to bridge disparate accounts of subiculum and investigate how subiculum neurons guide behavioral flexibility and navigation (i.e., speed, trajectory, and HD) have been observed in the subiculum (Kitanishi, Umaba, and Mizuseki 2021).

Subiculum neurons exhibit neural representations related to environmental geometry, such as place, corners, and boundaries (Brotons-Mas,et al. 2017; Kim et al. 2012; Lever et al. 2009; Sun et al. 2024). Our model currently does not account for all neural correlates in the subiculum, such as corner cells (Sun et al. 2024). However, the temporal dynamics of corner cells (once formed) could be accounted for in the present model. When the two walls forming a corner are gradually separated, corner cells also exhibit trace fields, which might be explained by our mismatch learning rule. Future work could investigate the trace fields of corner cells based on our proposed framework. As for generating corner cells, one possibility is to combine cells detecting walls on the left and right of the virtual rat facilitated by egocentric BVCs identified in the lateral entorhinal cortex (Wang et al. 2018).

## 5 Conclusions

Our work provides the first neural, circuit-level model of intra-subiculum computational mechanisms and CA1 interactions, combined with proximodistal differences in subiculum and CA1. As the proximal-distal CA1-subiculum pathway has been hypothesized in spatial memory specialization (Igarashi et al. 2014; Matsumoto et al. 2019; Nagelhus et al. 2023; Vandrey et al. 2021), future work could use our proposed framework to explore the neural mechanism of object and spatial information processing. This would deepen our understanding of the CA1-subiculum pathway and help integrate the subiculum with the canonical hippocampus model. Updating the cognitive map via the subiculum in accordance with recent experiences may also influence model-based reinforcement learning, where the hippocampal map serves as a world-model (Chersi and Burgess 2015). Extrapolating further, hippocampal involvement in non-spatial computations (Bicanski and Burgess 2019; Bottini and Doeller 2020; Constantinescu, O’Reilly and Behrens, 2016; Epstein et al. 2017; Theves, Fernández and Doeller 2020; Theves et al. 2021) might suggest a generalized role for the subiculum in updating all cognitive maps, including abstract maps (Behrens et al., 2018; Constantinescu, O’Reilly and Behrens 2016; Garvert, Dolan, & Behrens, 2017). Thus, our model may constitute a universal framework that could be employed for the updating during spatial and non-spatial cognitive mapping tasks alike.

## Acknowledgements

AB and FW acknowledge funding from the Max-Planck Society. We are grateful for several productive discussions with Colin Lever.

---

**Table 1:** Model parameters.

| Parameter | Description | Default |
| --- | --- | --- |
| $y_{cue}$ | The y-coordinate of the inserted cue | 0.5m |
| $N_{pSub}$ | Number of pSub neurons | 120 |
| $\delta\theta$ | Size of angular integration step for pSub BVCs | $2^\circ$ |
| $\xi$ | Inverse gradient in the linear function of tuning distance of pSub BVCs | 12 |
| $\sigma_0$ | Intercept in the linear function of tuning distance of pSub BVCs | 0.08 |
| $N_{CA1}$ | Number of CA1 neurons | 100 |
| $\sigma$ | The width of the CA1 place field | 0.05m |
| $w_{ij}(0)$ | The initial value of the CA1-dSub weights | 0 |
| $a$ | The hyperparameter of the sigmoid function | 1 |
| $b$ | The hyperparameter of the sigmoid function | 58.89 |
| $c$ | The hyperparameter of the sigmoid function | 0.1 |
| $\tau$ | The updating interval of the CA1-subiculum weights | 100ms |
| $N_\alpha$ | Number of distinct learning rates | 4 |
| $m_1$ | The parameter of the nonlinear function $Q$ | 3 |
| $m_2$ | The parameter of the nonlinear function $Q$ | 10 |
| $dx$ | The sampling interval along the inserted cue | 0.01m |
| $A_w$ | The maximum efficacy of the CA1-subiculum weights in models with cue-dependent modulation | 0.5 |

## Notes

### Competing Interest Statement

The authors have declared no competing interest.

